# NMR microsystem for label-free characterization of 3D nanoliter microtissues

**DOI:** 10.1101/2020.04.26.062141

**Authors:** Marco Grisi, Gaurasundar M. Conley, Kyle J. Rodriguez, Erika Riva, Lukas Egli, Wolfgang Moritz, Jan Lichtenberg, Jürgen Brugger, Giovanni Boero

**Affiliations:** Annaida Technologies SA, Lausanne, Switzerland; Microsystems Laboratory, École Polytechnique Fédérale de Lausanne (EPFL), Lausanne, Switzerland; Service de Gastro-Entérologie et d’hépatologie, CHUV, Epalinges, Switzerland; InSphero AG, Schlieren, Switzerland

**Keywords:** NAFLD, CMOS, NMR, nanoliter, microtissue, 3D cell culture, label-free, lipids, metabolism

## Abstract

Performing chemical analysis at the nanoliter (nL) scale is of paramount importance for medicine, drug development, toxicology, and research. Despite the numerous methodologies available, a tool for obtaining chemical information non-invasively is still missing at this scale. Observer effects, sample destruction and complex preparatory procedures remain a necessary compromise^1^. Among non-invasive spectroscopic techniques, one able to provide holistic and highly resolved chemical information *in-vivo* is nuclear magnetic resonance (NMR)^2,3^. For its renowned informative power and ability to foster discoveries and life-saving applications^4,5^, efficient NMR at microscopic scales is highly sought after^6–10^, but so far technical limitations could not match the stringent necessities of microbiology, such as biocompatible handling, ease of use, and high throughput. Here we introduce a novel microsystem, which combines CMOS technology with 3D microfabrication, enabling nL NMR as a platform tool for non-invasive spectroscopy of organoids, 3D cell cultures, and early stage embryos. In this study we show its application to microlivers models simulating non-alcoholic fatty liver disease (NAFLD), demonstrating detection of lipid metabolism dynamics in a time frame of 14 days based on 117 measurements of single 3D human liver microtissues.

## Introduction

Nuclear magnetic resonance (NMR) is a powerful spectroscopic tool for non-invasive investigations of the chemistry in intact living matter^2,3^. In the last decades, NMR has been a key enabler for several lifesaving diagnostic applications^4,5^ as well as groundbreaking fundamental research on animal and human bodies^11,12^. However, due to limitations of the current commercial hardware, magnetic resonance cannot be routinely applied to *in-vivo* and *in-vitro* studies at the nanoliter (nL) scale, typical of microorganisms and cell cultures. Indeed, micro-NMR is a longstanding technical challenge, and numerous researchers have proposed methods to achieve it, mainly using micro solenoids^6,7^, planar micro coils^8^, magic angle coil spinning (MACS) probes at ultra-high fields^9^, reaching sample volume sizes of about 10 nL. More recently, NMR spectroscopy of eggs of microorganisms having a volume of just 0.1 nL was reported in a relatively weak field strength of 7 T. Such advancement was possible thanks to a novel approach based on complementary metal-oxide semiconductor (CMOS) microchips that delivered high sensitivity, robustness, and a local and easily accessible sensing region^10^. All these works pushed NMR to the nL range, but sample handling issues and a high level of complexity remained an obstacle to high throughput studies with sufficient statistics, essential for biology, preventing widespread adoption. In this work we present an innovative device that combines the advantages of CMOS-based NMR probes with an unprecedented ease of use, allowing investigations of nL biological entities with a satisfactory statistic for the first time.

To show these capabilities, we here describe a first study using nL NMR on 3D microtissues (MTs), microscopic cell aggregates that mimic features and functionalities of their macroscopic counterparts. MTs have been introduced as a mean to simulate and probe disease conditions by allowing high throughput drug screening without need for biopsies of sick individuals, to better understand the mechanisms by which diseases operate and progress, and to reduce the need for animal testing, all of which could contribute to an accelerated development of new therapeutic treatments^13–16^. In this study we focus on human liver MTs (hLiMTs), specifically treated to simulate non-alcoholic fatty liver disease (NAFLD)^14,15^. NAFLD has become a global medical issue for 10-40% of adults from different demographics worldwide, and the mechanisms by which this disease progresses still eludes researchers^17,18^. This progressive disease is characterized by the initial accumulation of lipids in hepatocytes, predominantly in the form of triglycerides. The homeostasis of these hepatic lipids depends greatly on several pathways such as fatty acid uptake, *de novo* synthesis, oxidation, and very low-density lipoprotein (VLDL) secretion^19,20^. The regulation of these pathways and their metabolic profiles are fundamental to the progression of NAFLD, therefore, many researchers have focused their efforts in proteomic and lipidomic studies^21,22^. While these studies have been instrumental in understanding NAFLD, the techniques used rely on invasive methods, such as biopsies, and destructive analyses. The situation is analogous for MTs, for which an informative, quantitative and non-invasive technique, capable of characterizing individual specimens, is still missing^1,13^. In this study we successfully utilize NMR spectroscopy on single liver MTs and demonstrate the observation of a dynamic fatty acid metabolism associated to a diet switch inducing recovery from a steatotic condition.

## Results and Discussion

In our experiments we use hLiMTs generated *in-vitro* from human hepatic cells of healthy donors (3D Insight™ Human Liver Microtissues, InSphero AG, Switzerland)^14,15^. MTs are then exposed to different dietary regimes: a fasting diet is used to culture the livers in a lean state (as control), while a diabetic (fatty) diet induces steatosis (see Methods). The images (Fig. 1a-b) show a comparison between a lean and a steatotic micro-liver, with lipids stained in green. Clearly, the steatotic MT is characterized by an accumulation of lipids, main and typical manifestation of NAFLD^17,18^. In our study a dynamic metabolism is induced by a diet switch: we start at Day1 with a steatotic and a lean experimental group, and we create a third group by administering a fasting diet to steatotic livers for 14 days (Fig. 1c). The three resulting experimental groups are LEAN (healthy livers in fasting medium), SOD (steatotic livers “on diet”, i.e. in fasting medium), SOF (steatotic livers “on fat”, i.e. in diabetic medium). During the course of the experiments, media are routinely refreshed to maintain MT viability, and small batches of MTs are collected and analyzed in 6 time points as indicated in Fig. 1d (see Methods for experimental protocols).

**Figure 1:**
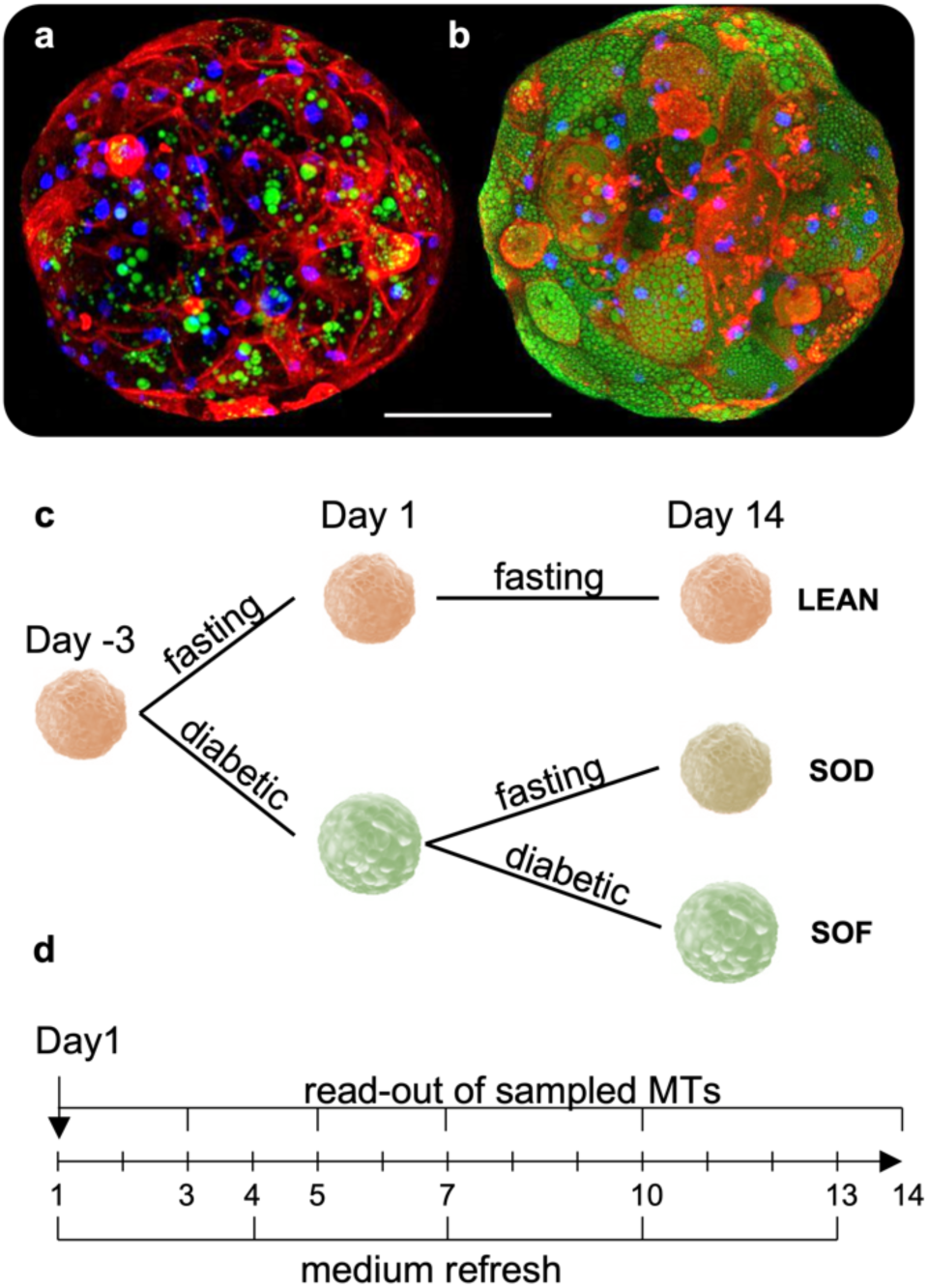
Samples and experimental design. Representative confocal images of human liver microtissues generated *in-vitro* after incubation in medium containing bovine serum albumin (BSA) **(a)** or oleic acid coupled to BSA (OA/BSA) **(b)** for 7 days. MTs were fixed and stained with Hoechst (nuclei, blue), CellMask™ (cell membrane, red) and Nile Red (lipids, green). The images clearly distinguish between a lean and fatty liver phenotype reminiscent of microvesicular steatosis. Scalebar = 100 μm. **(c)** Scheme depicting culture conditions of liver microtissues, starting after spheroid formation (day −3) and incubation until day 14 days with different media compositions, i.e. fasting medium (low insulin, low glucose) and diabetic medium (high insulin, high glucose) with fractionated plasma LDL. The three resulting experimental groups are lean livers (LEAN), steatotic livers “on diet” (SOD), steatotic livers “on fat” (SOF). **(d)** Read-out and medium refreshing time points.

Fig. 2 shows photographs and schematics of the CMOS-based NMR sensor (Annaida Technologies SA, Switzerland), consisting of a microsystem enclosed in a 3D printed plastic container (Veroclear, Connex500, Computer Aided Technology) holding the culture medium. The container has an openable lid, which presses on an o-ring (black in Fig. 2a) to seal the chamber and limit medium evaporation during measurements when closed, and allowing for quick sample access when open. The sensor connector is plugged into a dedicated receptacle, containing an electronic interface to the external console, prior insertion in the magnetic field (see S1). In this configuration, the sensor can be used as a plug & play device for an eased loading procedure.

**Figure 2:**
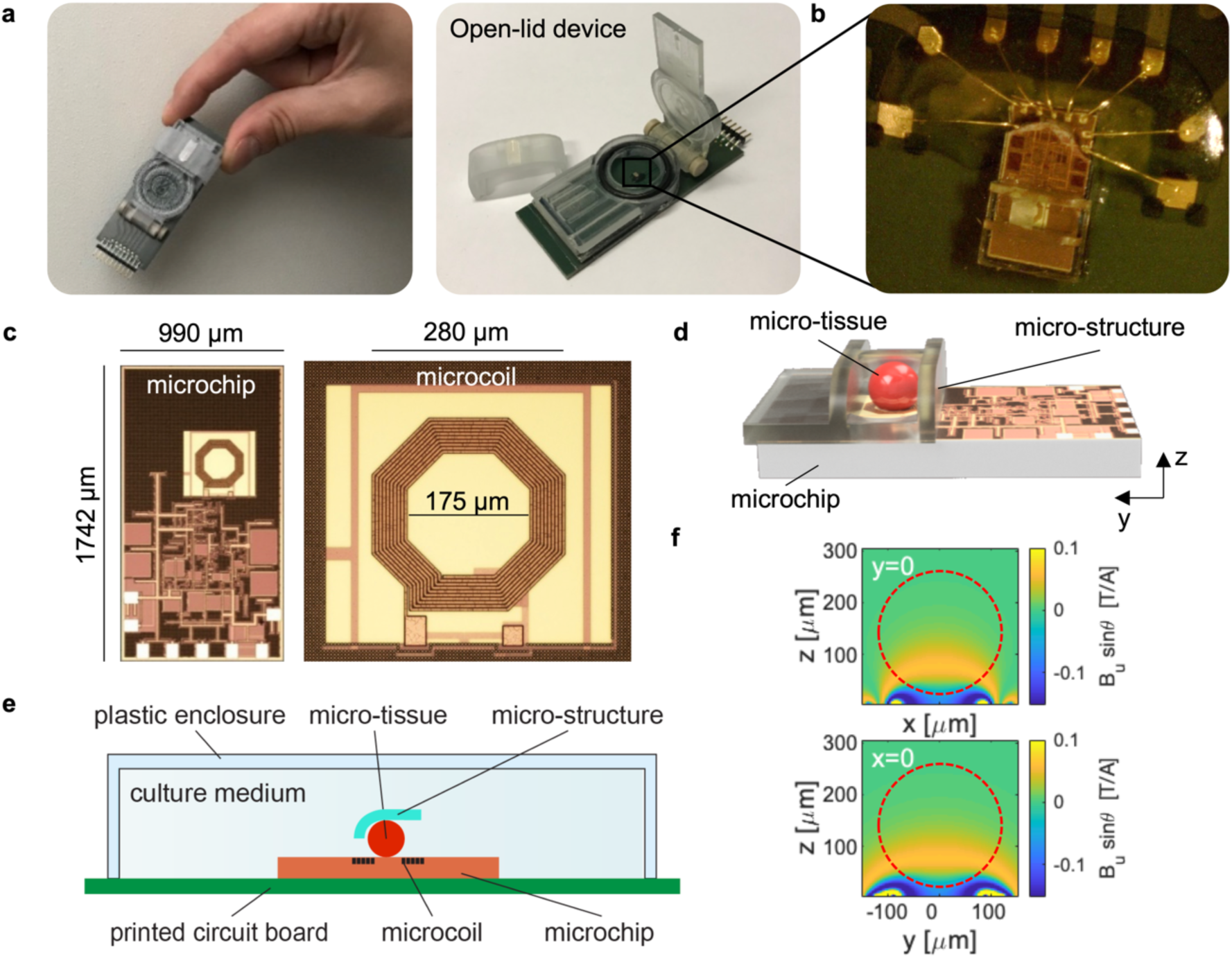
CMOS-based NMR probes for nL 3D MTs. **(a)** Photographs of our CMOS-based NMR sensor (6 x 2.5 cm) designed for in-vivo/in-vitro experiments at the nL scale. Left: sensor closed and ready for being inserted in the magnetic field. Right: sensor with open lid. **(b)** Photograph of the micro-system used to perform NMR spectroscopy of MTs. **(c)** Photographs of the CMOS microchip and its microcoil. The coil region appears with a different color due to the absence of metal density filling in order to minimize parasitic effects. **(d)** 3D rendering at scale of the micro-system and sample. The sample is approximated by a sphere having a diameter of 240 μm. In the indicated reference frame, the static field B_0_ and gravity are along the 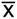 axis. **(e)** Cross-section schematics of the micro-system, defined as crossing transversally the chip at the microcoil center. The curved shape of the sample-positioning micro-structure enables placement of MTs of different sizes in the most sensitive area of the microcoil. **(f)** Maps of sensitivity of the integrated microcoil in experimental conditions (*τ*=20 μs, *i*=11 mA) at y=0 and x=0 cross-sections. The static *B*_0_ field is oriented along the 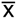 axis, while the 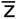 axis is perpendicular to the coil surface. The octagonal loops of the coil are approximated with circular ones. The unitary field *B*_u_, defined as the field produced by a current of 1 A in the coil loops, is computed via Biot-Savart. The component orthogonal to *B*_0_ (i.e. B_uzy_) is then considered to compute the nutation angle θ. Red dashed circular shapes indicate the sample as in (d-e). The excitation current *i*=11 mA is deduced by matching maximum signal intensity from experiments and sensitivity maps computation. This is in line with electronic simulations of the CMOS circuit in Cadence.

The microsystem (Fig. 2b) is made by combining a CMOS chip (Fig. 2c) with a 3D micro-structure made with a two-photon polymerization 3D printing technique (Photonic Professional GT by Nanoscribe GmbH, DE). This method, having a resolution below 1 μm^3^, allows fabricating structures that can easily be adapted to specific sample sizes and shapes. In this work the structure was specifically designed to position and hold MTs that are well approximated by spherical shapes (Fig. 1a-b). Our group had previously used this same fabrication technique to realize closed micro-channels for *in-vivo* measurements^23^. However, this previous system required a laborious mounting procedure, preventing the systematic accumulation of large, statistically relevant data sets. In the current device, a user-friendly design enables rapid sample placement for higher throughput experiments. The CMOS chip is wire bonded to the printed circuit board and the wires are covered with a protective glue. The sample-positioning micro-structure is then permanently attached to the chip surface to create a hosting spot for the MTs aligned on top of the integrated microcoil (Fig. 2d-e). Such pre-mounted micro-system is robust and allows for quick sample loading (< 5 minutes) with the sole use of a micro-pipette and a stereomicroscope. The whole set of measurements presented in this work was obtained by using only two identical sensors (see Methods).

Fig. 2f shows the sensitivity maps of the device as computed in the experimental conditions with a finite element model. Owing to the relatively long pulse used in the experiments (*τ*=20 μs, i=11 mA) a negative phase is originated close to the chip surface, which results in a reduction of the contribution to the total signal from the portion of medium close to the coil loops and outside the sample region by cancellation effects. In this configuration, the central part of the coil (i.e., where the sample is placed) has mainly a positive phase contribution, which constructively adds up in the total signal. From complementary experiments, we observed that the volume of the sample varies over time (see S2). Since in our system only a portion of the sample is analyzed (orange in Fig. 2f), the relation between signal intensity and volume is not expected to be linear. However, with the use of sensitivity maps, it is possible to estimate how sample volume affects the NMR signal and correct the data by factoring it out (see Methods). This is needed only if absolute quantities are compared.

Fig. 3 shows spectroscopy results of single MTs obtained with our CMOS-based sensor in a field strength of 7 T. Typical spectra are shown for acquisition times of 1 hour (Fig. 3a) and 10 minutes (Fig. 3b). The spectral resolution is about 0.2 ppm, slightly better than what observed by *in-vivo* liver studies^24^, and pronounced peaks are visible in the region between 0 and 6 ppm. In order to acquire a sufficiently large set of data to infer statistically significant conclusions, we fixed an experimental time of 10 minutes per single MT and performed several measurements of single MTs in the time points indicated in Fig. 1d. Figure 3c shows the 21 measurements acquired at Day14 for the three experimental groups, while the complete collection of the 117 spectra of single MTs is reported in S3. From averaged spectra at Day1 (Fig. 3d) and Day14 (Fig. 3e) it is evident that the SOD evolves to differentiate from SOF, expressing a phenotype more similar to LEAN in the long term. In the following, we interpret some of the peaks in our spectra to define numerical quantities (biomarkers) that can be used to follow this dynamic.

**Figure 3:**
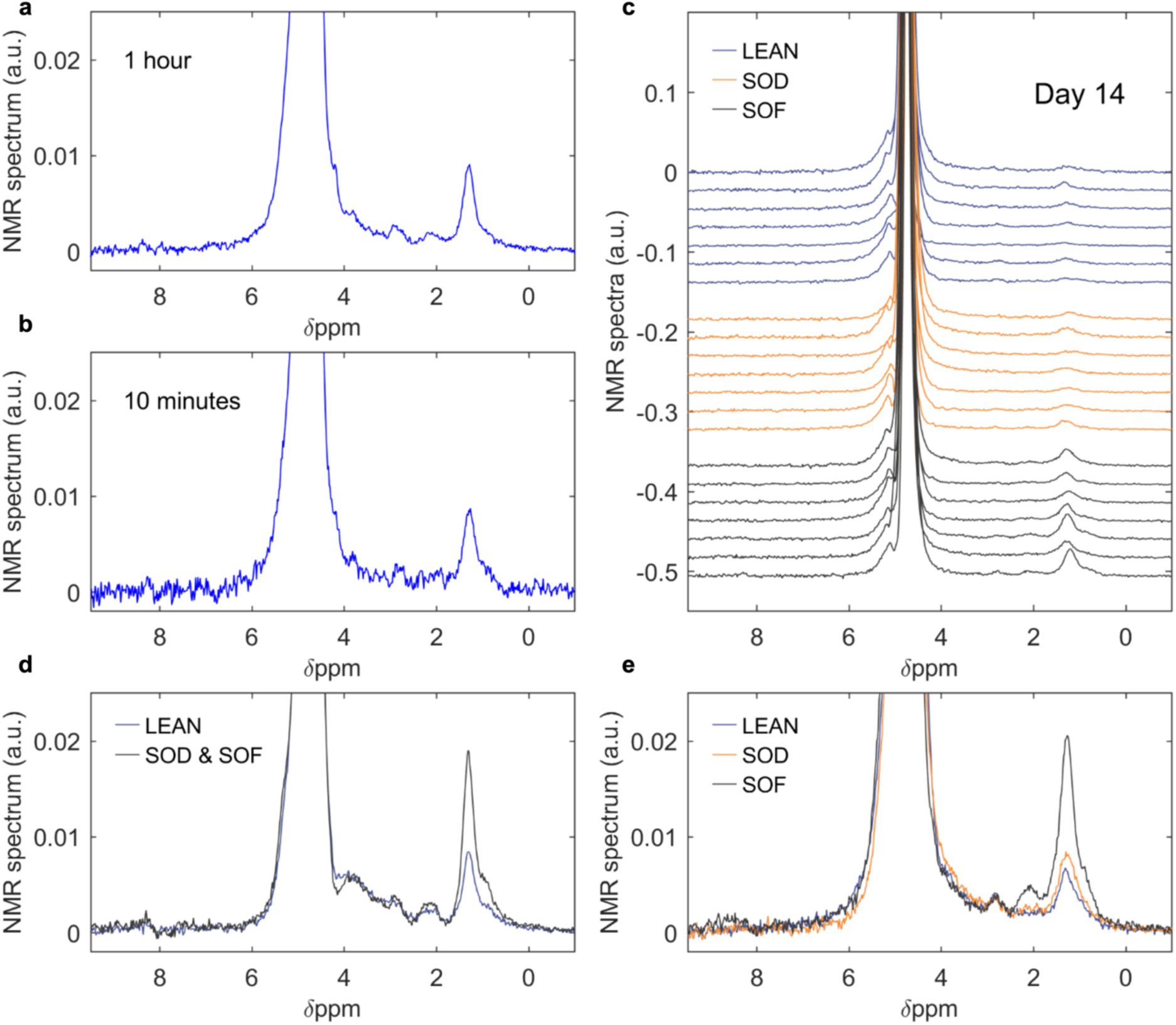
NMR spectroscopy of 3D microtissue cultures. **(a)** 1D ^1^H NMR spectrum obtained from a steaotic MT exposed to fasting medium (SOD) at Day10 over a measurement time of 1 hour, i.e. averaging 1800 scans. **(b)** 1D ^1^H NMR spectrum obtained from the same SOD MT as in (a) over a measurement time of 10 minutes, i.e. averaging 300 scans. **(c)** The complete series of measurements of individual MTs at Day14, where 7 measurements were collected for SOD, LEAN, and SOF MTs. Each measurement is obtained in 10 minutes time (i.e., averaging 300 scans). **(d-e)** NMR spectra averaged by experimental group at Day1 (d) and Day14 (e).

Previous literature reports have combined the use of ^1^H NMR spectroscopy on intact liver tissues and on extracts to determine the speciation of metabolites^25^ and to assess the role of lipids in liver quality^26^. Further evidence from *in-vivo* MRS of rat liver^24^ and NMR spectroscopy of oils^27^ suggests that the prominent signals shown in Fig. 3 correspond predominantly to lipids. With the scope of identifying efficient biomarkers, we restrict the analysis only to those signals that resonate sufficiently far from water to avoid overlap with its baseline. As discussed in S4-5, background and MT measurements indicate that the region ranging from 0 to 3.2 ppm can be considered for our purpose. Since metabolites are expected, in this spectral region, to give signals with a signal-to-noise ratio (SNR) lower by about one order of magnitude with respect to lipids^25^, we can consider their contribution negligible. To further corroborate this observation, lipidomic data obtained via mass spectroscopy of MTs extracts confirm that lipids are present in concentrations ranging from 0.1 to 1 M (see S6), in agreement with the observed 0.5 M of —(CH_2_)_n_ deduced from the signal amplitude at 1.3 ppm. These lipids are predominantly in the forms of triglycerides (TAGs), cholesteryl ester derivatives (CEs), and free fatty acids (FAs). While it is difficult to determine exact speciation with our SNR and spectral resolution, it is still possible to ascertain the types of functional groups and deduce information related to the degree of unsaturation of the lipids in the hLiMTs^24,27^ (see Table S7). As previously reported, a reliable way to obtain numerical values for statistical evaluations is to integrate over defined spectral regions^26^. In our case we identify three spectral regions of interest ranging from 1.18-1.46 ppm (integral L_1_), 1.80-2.20 ppm (integral L_2_), and 2.50-3.00 ppm (integral L_3_), which respectively represent aliphatic chains corresponding to the presence of lipid signatures from fully saturated (L_1_), mono-(L_2_) and poly-(L_3_) unsaturated fatty acids (see S7-8).

Fig. 4 shows experimental results and statistical significance of four different biomarkers defined to interpret the biological state of NAFL hLiMTs. Three of these markers are defined as ratios of the integrals L_i_ and represent relative concentrations of the aforementioned molecular groups. It is worth to note that these quantities, since defined by ratios of integrals computed from spectra of the same MT, are independent from the sample volume. In order to measure the biomarkers variability due to instrumental error (originating from electronic noise) and systematic errors (originating from sample placement, probe positioning, etc…), 12 additional measurements were performed on two MTs (6 each). The result of this investigation, presented in S9, shows that the variability due to error sources is well below the one observed across the different experimental groups, indicating that the defined markers provide a reproducible and sufficiently precise measurement to represent each MT. Fig. 4a-c show a direct comparison between values at the first and last time points. The SOD, significantly different from LEAN at Day1, approaches it differentiating from SOF in the long term. Fig. 4e shows the time evolution of L_3_/L_1_ monitored at all time points, where we can see its dynamics evolving gradually, but with a stepwise behavior between Day7 and Day10. Such behavior is also true for L_2_/L_1_, while for L_3_/L_2_ a change is already visible between Day5 and Day7 (see S10). A fourth marker, in analogy with conventional MRI-PDFF (Proton Density Fat Fraction, today’s gold standard and Endpoint in NASH trials)^28^ compares the content of —(CH_2_)_n_ groups to those found in LEAN at Day1. This marker, named Proportional Lipid Content (i.e. PLC), is a quantitative measurement of the saturated protons and it is corrected for MT volume variations. Fig. 4d shows that also PLC can well represent a dynamic evolution of the SOD from a steatotic to a lean status. Fig. 4f shows the overall dynamics, where we can see a gradual approach over time of the SOD from a lipid accumulation phase to a lean condition. The SOF, on the other hand, shows saturation of lipid content, reaching a maximum value already by Day 5 and remaining constant after that. In fact, the SOF showed no improvement from its steatotic state, while SOD MTs exhibited recovery reaching a state similar to that of LEAN. The data also reveal that while the PLC signal of SOD decreases between Day 7 and Day 14 towards lean values, L3/L1 shows a significant increase during the same period. This demonstrates that MTs under a fasting regimen not only reduce fatty acid content, but also gain in polyunsaturated fatty acid (PUFA) to saturated fatty acid (SFA) ratio. Similar alterations in hepatic PUFA/SFA ratio have been described in NAFLD/NASH patients compared to healthy control^22^. In summary, these data suggest that MTs are capable of reproducing a behavior similar to that exhibited by livers *in-vivo*, where studies relying on biopsies have shown that NAFL patients accumulated 5-10x the amount of fatty acids with an altered fats composition^21^, and fasting diets reduced lipid accumulation^29^. Similarly, in this study, the LEAN exhibited far less lipid content than SOF livers, and SOD MTs recovered to a lean status when exposed to a fasting medium.

**Figure 4:**
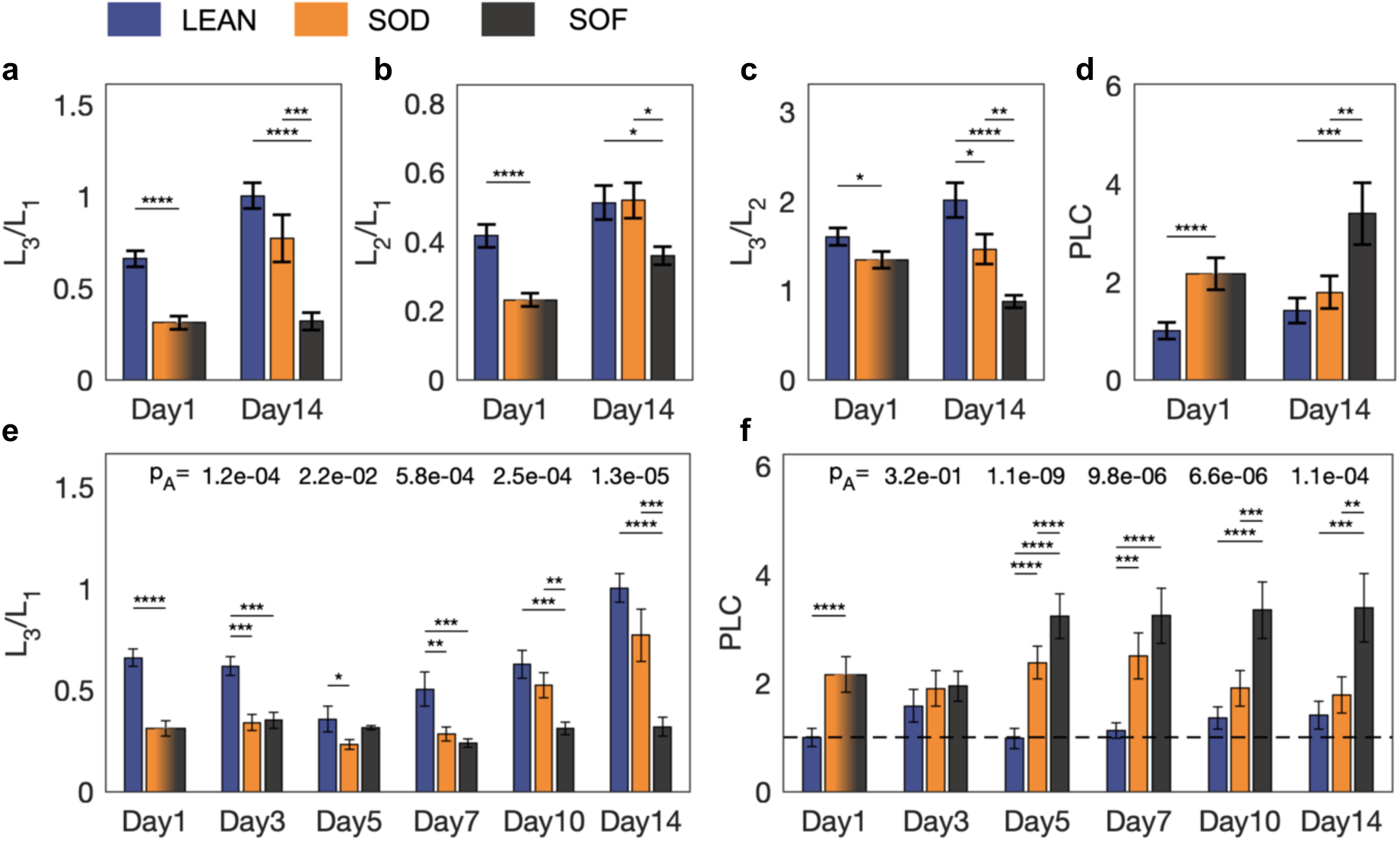
Label-free detection and evolution of lipids. Data are computed from spectra obtained by averaging 300 scans (i.e., 10 minutes measurement time). At each time point biomarkers of MTs are grouped as LEAN, SOD, SOF. At Day1, SOD and SOF coincide. 7 data points are acquired for SOD, except 5 data points at Day3. 7 data points are acquired for SOF, except 5 data points at Day14. For LEAN, 7 data points are acquired at Day1 and Day14, 6 data points at Day7 and Day10, 5 data points at Day3 and Day5. In total, 117 experiments are performed on single MTs. At Day1, significance is computed with a student t-test. From Day3, one-way ANOVA (p value ‘p_A_’ is indicated) and Tukey-Kramer tests are applied (biomarkers distributions are shown in S11). *P<0.05, **P<0.01, ***P<0.001, ****P<0.0001. L_1_: integral area from 1.18 to 1.46 ppm. L_2_: integral area from 1.8 to 2.2 ppm. L_3_: integral area from 2.5 to 3 ppm. PLC: L_1_/L_1_LEAN_Day1_. PLC values are corrected for volume variation to represent concentration values (see Methods). Direct comparison at Day1 and Day14 of: **(a)** L_3_/L_1_ **(b)** L_2_/L_1_ **(c)** L_3_/L_2_ **(d)** PLC. Complete time evolution of: **(e)** L_3_/L_1_ **(f)** PLC.

## Conclusion

In this work we have demonstrated and validated a device concept that enables NMR on single biological entities at the nanoliter scale. Its simplicity of use, sensing performance, robustness and versatility makes this technology suitable for extended high throughput non-invasive investigations of intact microscopic biological samples. The use of this CMOS-based NMR device is demonstrated in an exemplary study, detecting fatty acid metabolism dynamics, label-free, from more than 100 measurements of single nL human liver microtissues. The flexibility of the fabrication methodology makes CMOS-based micro-NMR a promising platform for more widespread use of the technique in different research environments. Such an advancement would enable non-invasive chemical analysis of biological entities that are currently out of reach, opening to novel applications based on new characterization criteria for a multitude of microorganisms, organoids, microtissues, and even early stage embryos, with great potential impact in medicine, drug development, and fundamental biochemical research.

## Methods

Experiments and protocols were approved by the SV-biosecurity unit committee of the École Polytechnique Fédérale de Lausanne and carried out in accordance with the experimentation guidelines of the institution.

### NMR experimental details

NMR experiments were performed in the 54 mm room temperature bore of a Bruker 7.05 T (300 MHz) superconducting magnet. The electronic setup is noticeably similar to the one described in detail in Ref. [30]. In this realization, the RF sources, DAQ, and TTL pulses are generated by a modified Tecmag Scout console, whose GUI is used to manage the experimental parameters. Two different frequencies are used in transmit and receive mode with a resulting down-conversion at about 11 kHz. Transmit frequency is fixed at the Larmor frequency. All experiments were performed with a repetition time of 2 s, a pulse length of 20 μs, and an acquisition time of 200 ms with 8192 data points. Prior measurements at each time point the magnetic field is shimmed with a probe without sample (i.e., a sample of culture medium) and left untouched for the day. The achieved spectral resolution with the culture medium is about 0.05 ppm full-widht-at-half-maximum.

### NMR data analysis

During the automated data analysis, all spectra are processed with strictly the same algorithm. The time domain data were post-processed by applying an exponential filter with decay time of 100 ms prior FFT. A frequency window going from 5 to 16 kHz is considered for analysis. An automatic algorithm is used to phase the spectra by maximizing the sum of the real part. The frequency axis is transformed to ppm units, and the maximum value of the water peak is aligned at 4.75 ppm. After baseline correction, the biomarkers are computed as explained in S8 and standard statistical analysis is applied to the resulting values. Overall, this study is based on 117 spectra of single micro-tissues, plus 12 spectra obtained on two micro-tissues in order to measure experimental systematic and instrumental errors.

### On-chip microcoil details

The on-chip microcoil is realized using 4 of the 6 metal layers available in the CMOS technology. 10 loops with pitch of 5μm are used on the top metal (metal 6), and 12 loops with pitch of 10.6 μm are equally distributed in three layers (metal 5,4,3). The loops are connected in series. The internal diameter of the coil is 180 μm.

### Correction for volume variation

With the use of sensitivity maps in Fig. 2f, it is possible to estimate how sample volume affects the NMR signal and therefore correct data factoring out sample volume dependency. For instance: by integrating the sensing map over the sample region shown in Fig. 2f (dashed red shape) we can compute the total signal from a sample having a diameter of 240 μm. The same calculation over a sample volume having a diameter of 200 μm shows that the signal intensity decreases by about 25%. Therefore, if absolute quantities deduced from spectra are compared from these two situations, a correction factor of 0.75 should be taken into account. When absolute quantities are compared (PLC in Fig. 4), this procedure is used to correct for sample volume variations. The numerical correction factors for MTs on fasting and diabetic medium, deduced from S2 and Fig. 2f at the six time points in Fig. 1d, are respectively: C_fasting_=[1,0.98,0.96,0.94,0.82,0.6]; C_diabetic_=[1,0.98,0.96,0.94,0.84,0.74].

### Statistical Analysis

Student’s t-test, or one-way ANOVA and Tukey-Kramer tests were used to calculate significance levels between groups with a Matlab program. Graphs and figure legends are annotated with the level of significance between the test groups.

### *In vitro* generation of MTs and culture

All spheroid hLiMT used in this study were 3D InSight™ Human Liver Microtissues (InSphero AG, Schlieren, Switzerland) produced according to the patent-pending protocol (WO2015/158777A1), composed of primary human hepatocytes (PHH) and non-parenchymal liver cells (NPCs). PHHs (pooled fractions originating from 10 individual donors) and NPCs were purchased from BioIVT (Westbury, NY). Human Liver Microtissues were provided and cultured in InSphero’s Akura™ 96-well plate format. Steatosis was induced either by oleic acid (400 μM coupled to BSA at a molar ratio of 6:1) for 7 days or by medium supplementation of fractionated LDL (Lee Biolsolutions, MO, USA) for 4 days. 3D InSight™ Human Liver Microtissues, 3D InSight™ Human Liver Maintenance Medium (TOX) were obtained from InSphero AG, Schlieren, Switzerland.

### Culture media

According to the desired phenotype, the three experimental groups of liver PHHs and NPCs co-culture (3D microtissues produced by InSphero) were treated with two different culture media (Fig. 1c). These same media are used to obtain the starting (i.e. Day1) lean and steatotic phenotypes. The LEAN and SOD were kept in a physiological medium (fasting medium), based on William’s E, additionally containing glucose, insulin, glucagon, pyruvate and lactate, for 14 days, resulting in a lean phenotype. A second, more anabolic medium (diabetic medium) is used to obtain the steatotic phenotype, based on William’s E additionally containing glucose/fructose, insulin as well as plasma fractionated LDL. The SOF is cultured in this medium for 14 days but washed and measured within fasting medium to allow for a direct comparison among experimental groups (see protocol below). Medium exchange was performed simultaneously in each group every 2-3 days as indicated in Fig. 1d.

### Preparation of MTs for confocal imaging

MTs were transferred with a tip quickly coated in fetal calf serum (FCS) into a CellCarrier-384 Ultra Microplates (6057300, Perkin Elmer). They were then fixed in 4% PFA (P6148, Sigma-Aldrich) for 1 h at room temperature. After washing with PBS, MTs were stained with Nile Red for lipid droplets, Hoechst for nuclei and CellMask™ (C10046, ThermoFisher) for plasma membrane for 1h at room temperature on an orbital shaker. Subsequently, MTs were washed with and left in PBS until imaging.

### Confocal imaging

A Visitron spinning disc confocal microscope was used to acquire the images in Fig. 1a-b. In order to acquire the whole tissue, the focus was set to the bottom of the plate and the maximum z-range was imaged. Each acquired channel, its excitation/emission wavelength, and the corresponding exposure time are: (1) confDAPI, 405/461 nm, 250 ms; (2) confGFP, 488/525 nm, 250 ms; (3) confCy5, 640/666 nm, 250 ms

### Protocol for read-out of sample aliquots

Prior read-out, aliquots of 7 MTs are transferred from their original 96 well plate to a single Petri dish where they are immersed in the fasting medium. Such transfer is done by using a standard 200 μL pipette. In the case of the SOF, where the 96 well plate contains the MTs in diabetic medium, the aliquots are washed in fasting medium before being placed in the final Petri dish from which they are picked up for measurement (also containing fasting medium). For LEAN and SOD, since they are already immersed in fasting medium during culture, there is no washing step. After the transfer is done under a laminar flow, the culture plate is put back in the incubator and the Petri containing the aliquots is placed on a heated stage set at 37°C in laboratory atmosphere. From this point, samples are sequentially transferred from the Petri dish to the CMOS-based NMR sensors, which were previously filled with fasting medium. This second transfer is done by using a 300 μm pipette for embryology by Cook Medical and with the help of a stereomicroscope. Once a MT is loaded into the sensor and the measurement starts, the successive MT is loaded into a second identical sensor and prepared for the next experiment. By alternating between two sensors it is possible to reduce the experimental time and therefore the exposure of MTs to the laboratory atmosphere. Overall, this procedure took less than 90 minutes for all aliquots analyzed. During read-out the three experimental groups LEAN, SOD, SOF are treated separately, sequentially, and in no particular order. Out of the 7 samples picked for the study, some were occasionally lost therefore reducing the experimental points to 6 or 5 in a few time points (see Fig. 4 legend for MTs number).

## Author Contributions

GMC, MG, JL, WM conceived the study. LE, WM, JL provided samples and protocols. MG and GMC designed and fabricated the CMOS-based sensors. MG and GMC performed experiments and data collection. GB, JB provided critical analysis and discussion. MG, KJR, ER performed data analysis. MG supervised the project. MG and KJR wrote the manuscript. All authors edited and approved the content.

## Acknowledgements

We thank Margaux Duchamp from LMIS4 laboratory (EPFL) for allowing use of cell culture incubators. We thank prof. Lyndon Emsley for fruitful discussions. We are grateful to the help of the staff at the Center of Micro and Nano Technology (CMi) at EPFL with the fabrication process. This work was partially supported by EPFL, the Swiss National Science Foundation under grant 40B1-0_180268, the FIT foundation of Canton Vaud via the EPFL Innogrant program, Innosuisse under grant 39821.1 IP-LS, and the European Union’s Horizon 2020 research and innovation program under grant agreement No 681002.

## Conflicts of interest

MG, GMC, JB, and GB are the authors of a patent application related to this work (Submitted on September 24^th^ 2018, PCT/IB2018/057348). GMC and MG are co-founders of Annaida Technologies, which owns an exclusive license of the mentioned patent application.

## Notes

### Competing Interest Statement

MG, GMC, JB, and GB are the authors of a patent application related to this work (Submitted on September 24th 2018, PCT/IB2018/057348). GMC and MG are co-founders of Annaida Technologies, which owns an exclusive license of the mentioned patent application.

### Summary of Updates

Supplementary figure S9 has been corrected.

