## Supplementary Information for "NMR microsystem for label-free characterization of 3D nanoliter microtissues"

### S1: MicroNMR probe and CMOS-based sensors

Fig. S1 shows photographs of the CMOS-based NMR probe currently installed in our laboratory. This system, based on 3D printed supports, is easily customizable for any magnet currently available in laboratories worldwide.

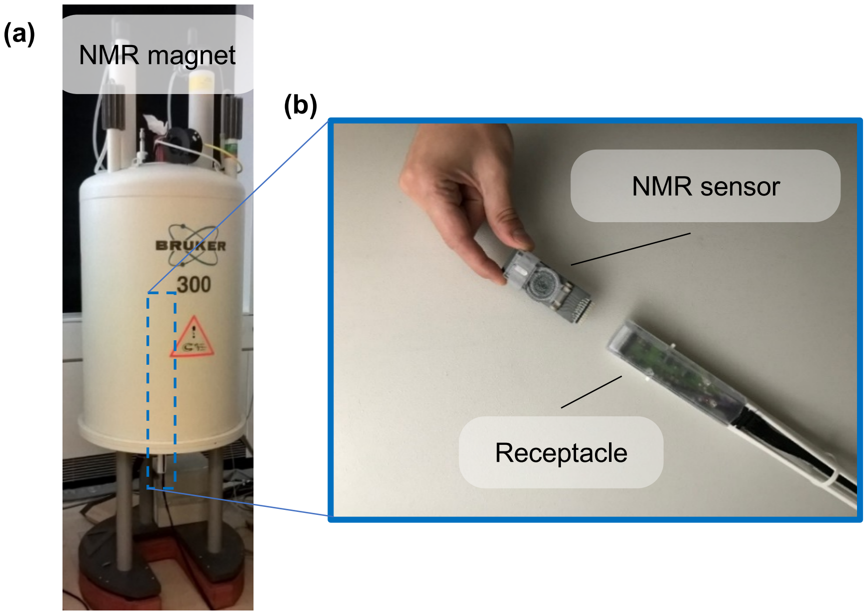


**Fig. S1. Prototype of CMOS-based NMR probe. (a)** Photograph of a vertical superconducting 300 MHz magnet (Bruker Biospin, Switzerland) with mounted probe including receptacle and sensor (Annaida Technologies SA, Switzerland). The NMR sensor can be easily inserted in the magnet, or retrieved, thanks to a 3D printed support that guides the probe into place via a slide & lock mechanism. **(b)** Photograph of sensor and receptacle. The micro-system is contained on a plug & play 6.5x2.5 cm board. For use the sensor is plugged into a receptacle and inserted into a magnetic field. Console and GUI complete the system.

### S2: Time evolution of MT volumes

To assess the variation of the sample volume over the experimental time, two groups of MTs have been exposed to a fasting and diabetic medium and their physical size has been measured with an optical microscope in three time points over a period of 15 days. MT sizes are reduced over time and exposure to different metabolic medium conditions. The results indicate that the MTs exposed to the diabetic medium reduce less in size, indicative of elevated fat accumulation. These data, combined with computations obtained with sensitivity maps, allow to correct absolute signal strengths for the natural sample volume variation. Such corrections are necessary only when absolute quantities at different time points are compared.

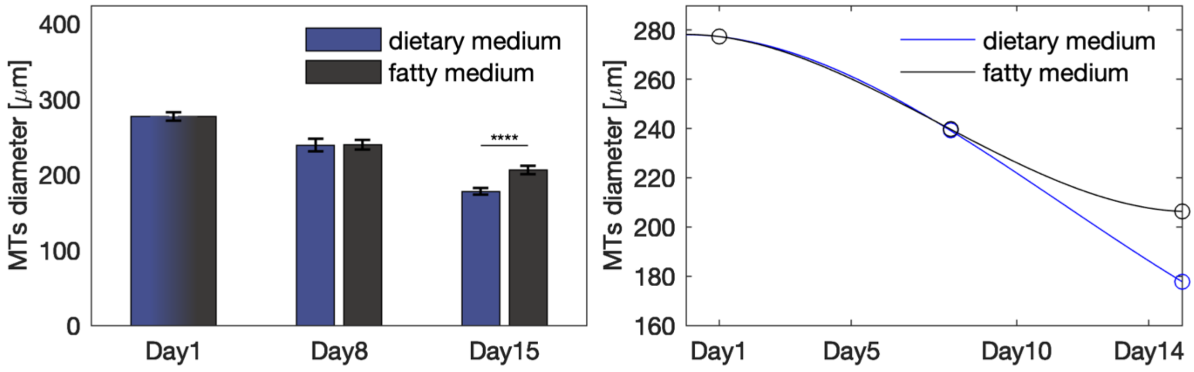


**Figure S2: Variation of MT volumes during culture.** At each time point MT diameters are measured and grouped according to the culture medium. 24 data points are acquired at Day1, 8 data points per group at Day8 and Day15. Significance levels are computed with student t-tests. *P<0.05, **P<0.01, ***P<0.001, ****P<0.0001. Left: Comparison of MT diameters. Right: Diameter values and polynomial fits used to interpolate at the experimental time points.

### S3: Complete series of MT measurements

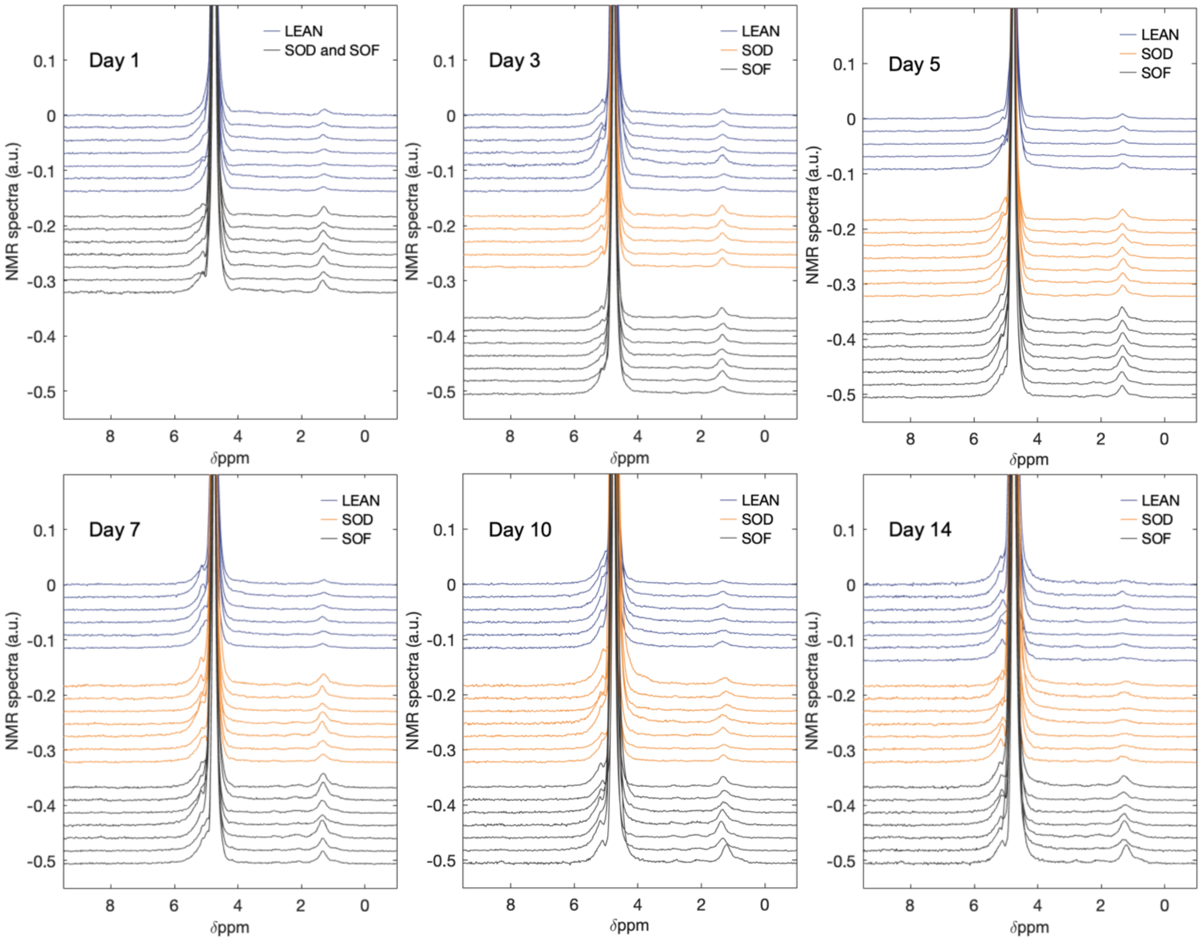


**Fig. S3. Complete collection of MT measurements.** 1D $^{1}H$ NMR spectra obtained from 117 single MTs in the 6 experimental time points as indicated in Fig. 1d. Each spectrum results from a measurement time of 10 minutes, i.e. averaging 300 scans.

### S4: NMR spectra of 3D micro-livers cultures

Fig. S4 shows the same spectrum of Fig. 3a-b plotted with full range along the y-axis. In the main graph we can see a prominent signal at about 5.2 ppm: due to the presence of a strong baseline from the water signal, the 5.2 ppm peak is excluded as a source to compute biomarkers in the methods developed in this work. As shown in the inset, some information is already visible after 10 minutes of averaging between 0 and 3.2 ppm.

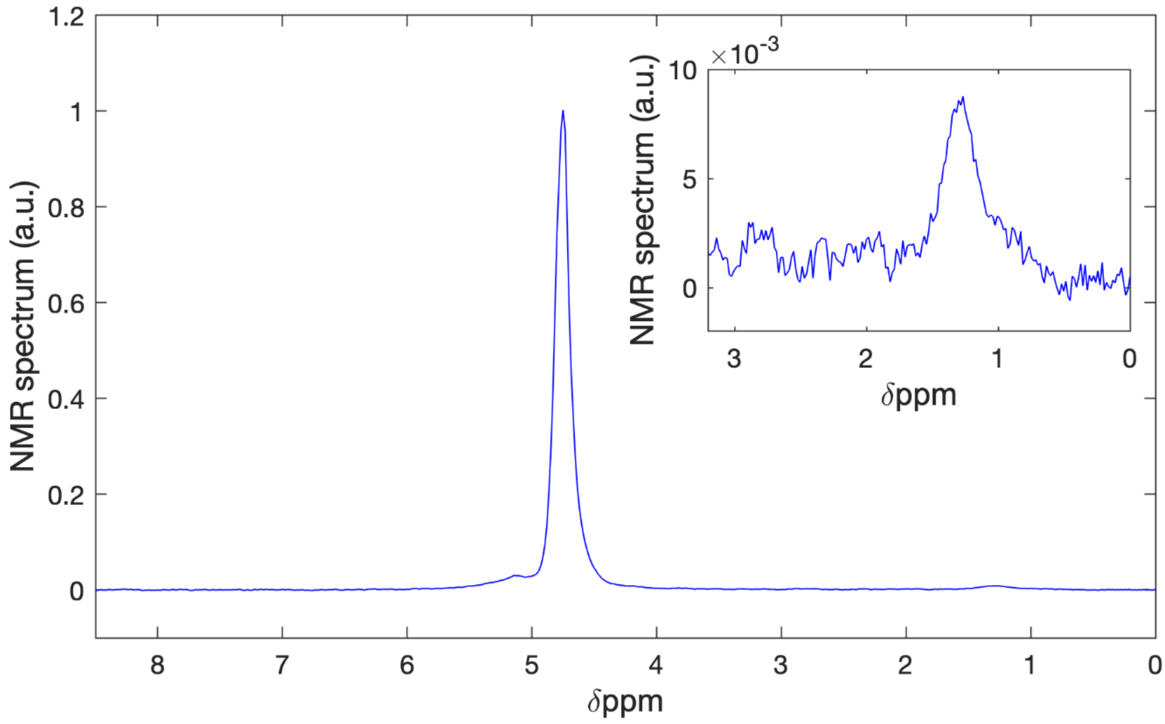


**Fig. S4. Spectroscopy of 3D micro-liver cell-cultures.** 1D $^1$H NMR spectrum obtained from the same steaotic MT as in Fig. 3a-b over a measurement time of 10 minutes, i.e. averaging 300 scans. The water signal is normalized to 1.

### S5: NMR Spectrum of fasting medium

The fasting medium was used to culture the LEAN and as the dietary medium for the SOD. Additionally, all NMR spectra recorded for this study were obtained from MTs immersed in this medium. In order to evaluate if the medium itself could lead to background signals in the region of interest for this study (i.e. from 0 to 3.2 ppm), Fig. S5 shows a direct NMR measurement. The resulting spectrum highlights that there are no visible signals in the region of interest.

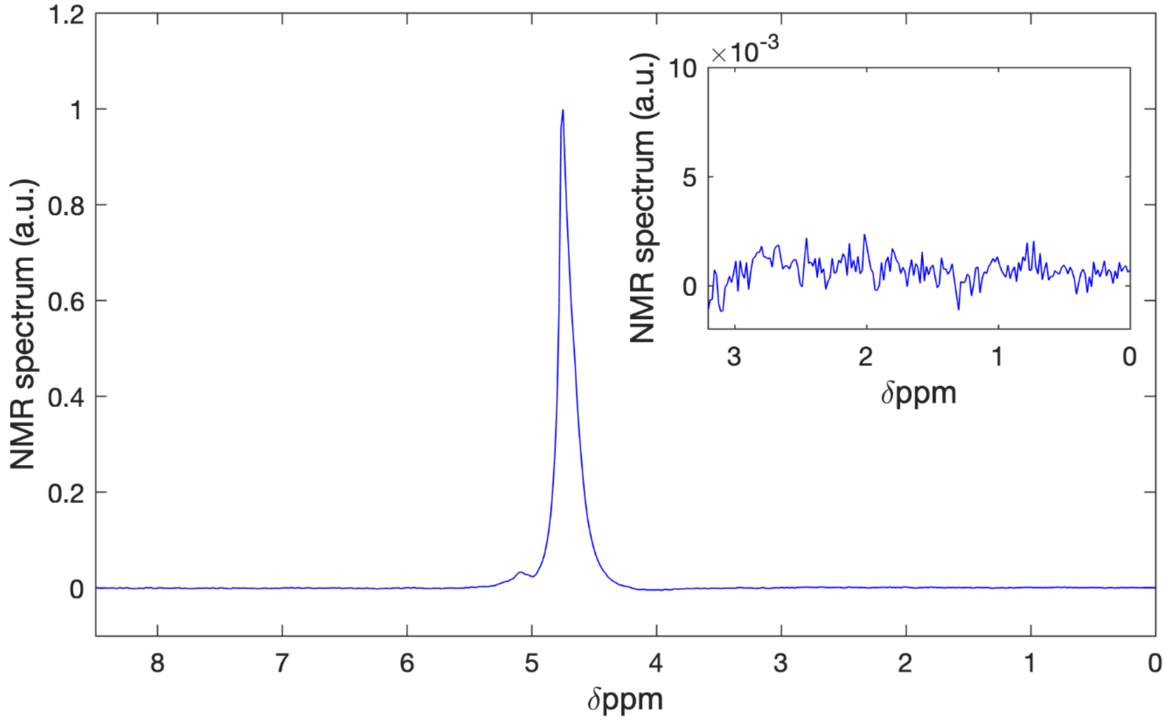


**Fig. S5. $^{1}$H NMR spectrum of fasting medium.** 1D $^{1}$H NMR spectrum obtained from the fasting medium over a measurement time of 10 minutes, i.e. averaging 300 scans. The inset illustrates that the region corresponding to the lipid resonances lacks any peaks.

### S6: Mass spectroscopy lipidomics of liver MTs

Microtissues were incubated for one week with either lean or diabetic medium to obtain two groups: lean MTs, and fatty MTs. The profile demonstrates the expected differences of detectable lipid classes, i.e. the increased appearance of Triglycerides (TGA), Cholesteryl ester (CE), Phosphatidylcholine (PC), free Cholesterin (Chol), Sphingomyelin (SM), Diacylglycerides (DAG), all components of LDL. Despite trends showing the expected separations, the large standard deviations and the low number of samples analyzed (3 samples per experimental group) result in poor statistical significance. Such a low statistics is due to: (1) the necessity to pool 15 MTs to obtain a single measurement; (2) the high experimental and economic costs of this analysis.

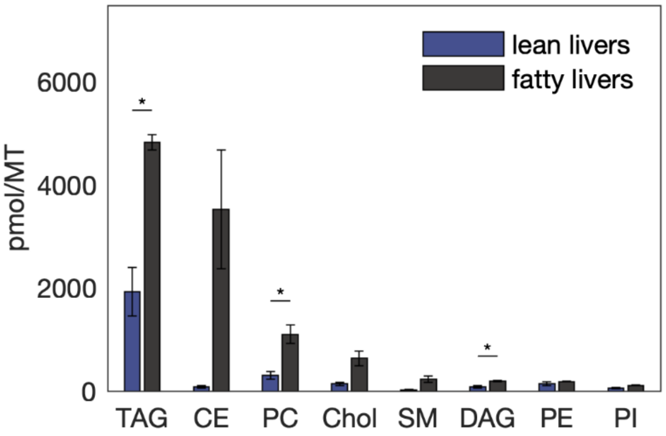


**Fig. S6. Lipidomics profiles of liver MTs with mass spectroscopy.** Shotgun lipidomics profile (by Lipotype, Germany) from microtissue lipid extracts ($MeOH/CHCl_3$), measured on a high-resolution Orbitrap mass spectrometry device (Thermo Scientific Q-Exactive). Significance levels are computed with student t-tests. *P<0.05, **P<0.01, ***P<0.001, ****P<0.0001.

### S7: Assignments of relevant chemical shifts in lipids and liver tissues

From literature reports of $^1$H NMR spectroscopy of liver tissues and extracts, as well as oils, it is possible to deduce the chemical shifts of all relevant functional groups in liver lipids. In Ref [1], where liver fat modulation was investigated in rats by $^1$H NMR, a spectrum is shown containing similar resonances as in the CMOS-based measurements performed in this work. The main resonances were attributed to methyl, methylene, and branched unsaturated hydrocarbons affiliated with lipids predominantly in the forms of triglycerides (TAGs), cholesteryl ester derivatives (CEs), and free fatty acids (FAs)[2,3]. NMR of extracts and oils further elucidate the features of lipids in $^1$H spectra[4-7]. Table S7 reports the chemical shifts shown with high resolution spectroscopy, where the different chemical groups and the respective chemical shifts are reported. Based on these data, we have defined 4 groups of interest that relate to saturated protons ($L_1$, δ=1.18-1.46 ppm), mono-unsaturated protons ($L_2$, δ=1.8-2.2 ppm), poly-unsaturated protons ($L_3$, δ=2.5-3.0 ppm), mono-unsaturated protons ($L_4$, δ=5-5.4 ppm). For our purpose, in this work we concentrate on the region going from 0 to 3.2 ppm in order to avoid any overlap with the baseline from the water signal. However, it is worth noting that looking further downfield at the spectra, resonances from roughly 3.8 ppm to 7 ppm are of interest and can further help to evaluate molecules affiliated with liver disease in literature.[6-8] The compounds that appear in this region are glycogen, cholesterol, CEs, PCs, LPCs, triacylglycerols, SMs, PEs, glucose, and the group $L_4$ identified below.

**Table S7. Lipids chemical shifts.** Chemical shift of lipids as reported in literature and separation into functional groups for MT spectra analysis. Chemical shifts are reported from Ref. [6].

| Chemical group | $^1$H Chemical Shift (δ) (Literature) | $^1$H Shift range (δ) | Group Name |
|---|---|---|---|
| **$CH_3$**— | 0.83-1.03 | / | / |
| —**($CH_2$)$_n$** | 1.22-1.42 | 1.18-1.46 | **$L_1$** |
| —**$CH_2$**—$CH_2$—$CO_2$— | 1.52-1.70 | / | / |
| —CH=CH—**$CH_2$**— | 1.94-2.14 | 1.80-2.20 | **$L_2$** |
| —**$CH_2$**—$CO_2$— | 2.23-2.36 | / | / |
| =CH—**$CH_2$**—CH= | 2.70-2.84 | 2.50-3.00 | **$L_3$** |
| —**CH**=**CH**— | 5.26-5.40 | 5.00-5.60 | $L_4$ |

In order to gain a better understanding of how these classes of compounds appear spectrally using the CMOS-based probe, free fatty acid (FFA) standards were chosen based off of their unsaturation and chain length. In this study, the FFAs investigated were palmitic acid ($C_{16}$, sat.), oleic acid ($C_{18}$, mono.), and linoleic acid ($C_{18}$, poly.). All standards were prepared in DMSO-$d_6$ at a concentration of 1M. Using this system, it is clear that each compound can be differentiated by its degree of unsaturation and chain length.

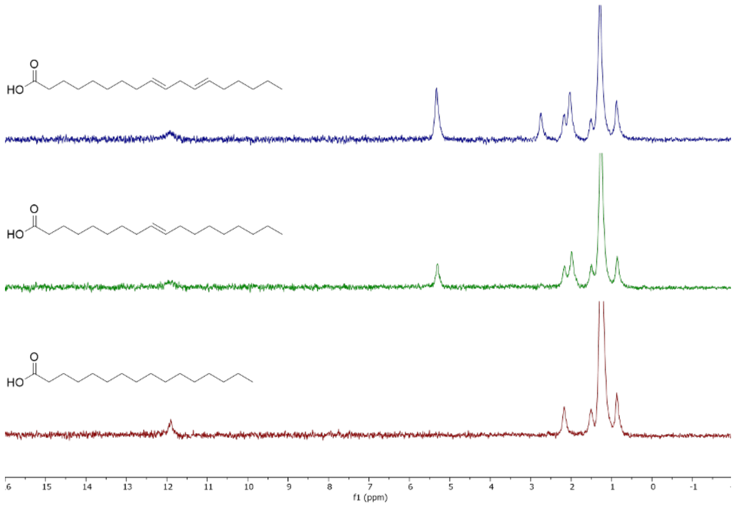


**Fig. S7. Stacked 1D $^1$H NMR spectra of Free Fatty Acid Standards.** Spectra of palmitic acid (red), oleic acid (green), and linoleic acid (blue) recorded using the same acquisition parameters used for the MT measurements, but averaging only 100 scans.

### S8: Computation of biomarkers from NMR spectra

In order to build a statistical analysis of the large number of measurements obtained in this study, it is necessary to define values associated to features of the measured spectra. In our case we observe spectra with three prominent peaks that are sufficiently far from the water signal so that their intensity is not affected by it: they relate to saturated protons (from 1.18 to 1.46 ppm), mono-unsaturated protons (from 1.8 to 2.2 ppm), poly-unsaturated protons (from 2.5 to 3 ppm). In Fig. S8 it is shown a spectrum with the associated integrals $L_1$, $L_2$, $L_3$. The ratios of these integral areas are defined as biomarkers as they relate to the chemical composition, in relative (and not absolute) terms, of each sample.

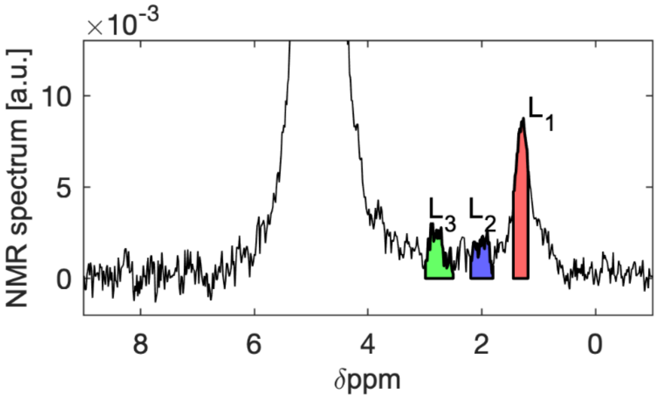


**Fig. S8. Biomarkers computation from NMR spectra.** 1D $^1$H NMR spectrum obtained from the same single steatotic MT as in Fig. 3a-b over a measurement time of 10 minutes, i.e. averaging 300 scans. In red: integral area $L_1$ of saturated protons (from 1.18 to 1.46 ppm). In blue: integral area $L_2$ of mono-unsaturated protons (from 1.8 ppm to 2.2 ppm). In green: integral area of poly-unsaturated protons (from 2.5 to 3.0 ppm).

### S9: Measurement of systematic and instrumental errors

In order to verify that the biomarkers L3/L1, L2/L1, L3/L2 computed from NMR spectra are representative of sample properties, we here demonstrate reproducibility as well as sufficient precision to distinguish different groups of samples. To elucidate these aspects, we have performed 12 experiments on two different MTs (MT1 and MT2 below) at Day10. 6 experiments were dedicated to estimate the instrument error: in this case we have observed MT1 acquiring 6 consecutive measurements without moving the sample. Since the electronic noise distribute randomly between separate scans, this allowed us to estimate the error on the biomarkers isolating the noise contribution from the instrument (values indicated by black asterisks). 6 experiments were dedicated to estimate the systematic errors: in this case we measured MT2 by alternating the sample between two identically designed sensors (same procedure used for the study, see Methods). This allowed us to estimate the error on the biomarkers associated to sample placement, sensors replication, and loading procedure (values indicated by blue squares). Fig. S9 below demonstrates the reproducibility and reliability of the read-out for all the three biomarkers here defined. We can also see that instrumental errors are of the same order of magnitude as systematic errors associated to manipulation, sample placing, sensors replication.

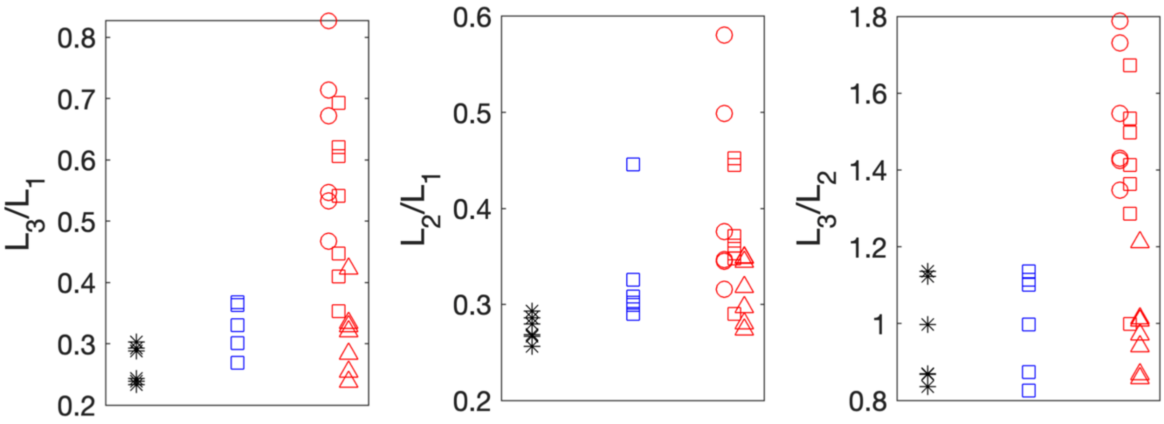


**Fig. S9. Direct measurement of instrument precision and systematic errors.** Measurements are obtained at Day10 from two steatotic micro-tissues (one from SOF, another from SOD) and L3/L1, L2/L1, L3/L2 are evaluated from measurements obtained by averaging 300 scans (i.e., 10 minutes measurement time). In black (stars) is plotted the result from 6 measurements where MT1 (SOF group) is left in position inside the probe. In blue (squares) is plotted the result from 6 measurements where the MT2 (SOD group) is each time transferred and re-positioned. In red are plotted all the results obtained from the MTs analyzed at Day10 (MT1 and MT2 are excluded from the series), circles refer to LEAN, squares to SOD, triangles to SOF. Overall, these data are based on 32 spectra obtained from 22 different MTs. The resulting values from MT1 and MT2, compared to all data at Day10, demonstrate the reproducibility and reliability of the read-out and give estimations for both the instrument precision as well as systematic errors.

### S10: Time evolution of biomarkers

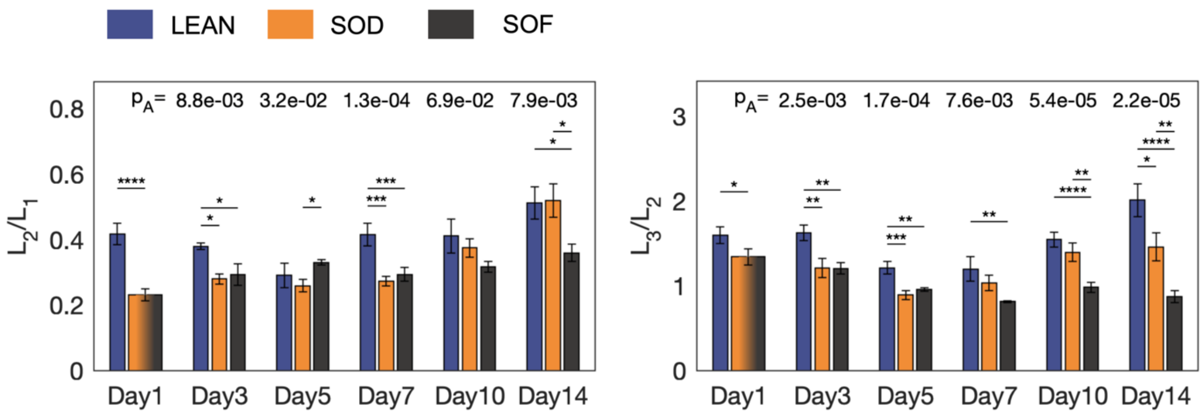


**Figure S10: Label-free detection and evolution of lipids.** Data are computed from spectra obtained by averaging 300 scans (i.e., 10 minutes measurement time). At each time point biomarkers of MTs are grouped as LEAN, SOD, SOF. 7 data points are acquired for SOD, except 5 data points at Day3. 7 data points are acquired for SOF, except 5 data points at Day14. For LEAN, 7 data points are acquired at Day1 and Day14, 6 data points at Day7 and Day10, 5 data points at Day3 and Day5. In total, 117 experiments are performed on single MTs. At Day1, significance is computed with a student t-test. From Day3, one-way ANOVA (p value '$p_A$' is indicated) and Tukey-Kramer tests are applied. *$P<0.05$, **$P<0.01$, ***$P<0.001$, ****$P<0.0001$. $L_1$: integral area from 1.18 to 1.46 ppm. $L_2$: integral area from 1.8 to 2.2 ppm. $L_3$: integral area from 2.5 to 3 ppm. Left: Complete time evolution of $L_2/L_1$. Right: Complete time evolution of $L_3/L_2$.

### S11: Distributions of Biomarker values

Evaluation of statistical difference between grouped data is made with standard tests (Fig. 4). These statistical tests make the assumption that each data point distributes randomly following a normal distribution with the same variance. Fig. S11 shows the distributions of the biomarkers measured experimentally in this work. Since variances are similar, deviations are obtained by subtracting the mean value within each LEAN, SOD, SOF experimental group and all 117 values are pooled. Distributions are obtained distributing uniformly 12 bins. Overall, Fig. S11 shows that our set of biomarkers follow normal distributions in good approximation.

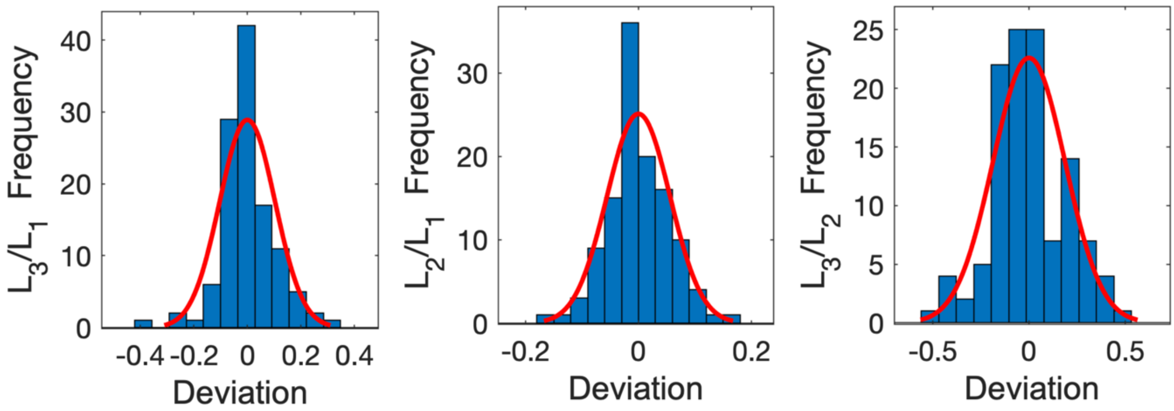


**Fig. S11. Distribution of Biomarkers.** Distributions of L3/L1, L2/L1, L3/L2 are fitted to the nearest normal distribution.
